# Protein-protein binding pathways and calculations of rate constants using fully continuous explicit solvent simulations

**DOI:** 10.1101/453985

**Authors:** Ali S. Saglam, Lillian T. Chong

**Affiliations:** Department of Chemistry, University of Pittsburgh, Pittsburgh, PA 15260

## Abstract

A grand challenge in the field of biophysics has been the complete characterization of protein-protein binding processes at atomic resolution. This characterization requires the direct simulation of binding pathways starting from the initial unbound state and proceeding through states that are too transient to be captured by experiment. Here we applied the weighted ensemble path sampling strategy to enable atomistic simulation of protein-protein binding pathways. Our simulation generated 203 fully continuous binding pathways for the bacterial proteins, barnase and barstar, yielding a computed k_on_ that is within error of experiment. Results reveal that the formation of the “encounter complex” intermediate is rate limiting with ~11% of all diffusional collisions being productive. Consistent with experiment, our simulations identify R59 as the most kinetically important barnase residue for the binding process. Furthermore, protein desolvation occurs late in the binding process during the rearrangement of the encounter complex to the native complex. Notably, the positions of interfacial crystallographic water molecules that bridge hydrogen bonds between barnase and barstar are occupied upon formation of the native complex in our simulations. Our simulations were completed within a month using 1600 CPU cores at a time, demonstrating that it is now practical to carry out atomistic simulations of protein-protein binding processes, particularly using the latest GPU-accelerated computing.

## 1. Introduction

Life is enabled by protein-protein interactions. Each of these interactions involves an elaborate molecular dance of the partner proteins until they fit together like pieces of a jigsaw puzzle. The steps of this dance – the binding mechanism – have been elusive to laboratory experiments due to the difficulty of capturing transient states. Furthermore, the binding mechanism can only be understood by analyzing pathways. Simply put, pathways *are* the mechanism.

Molecular dynamics (MD) simulations can generate pathways at the atomic level with high temporal resolution. However, due to the computationally prohibitive timescales of protein-protein binding processes, a limited number of studies have reported such simulations of protein-peptide binding^1–5^ and only one study has carried out such simulations of protein-protein binding.^6^ In the latter study, a Markov state model was constructed from a combination of independent trajectories and adaptive sampling to characterize the rapid binding kinetics of the proteins, barnase and barstar.^6^ While this study reproduced the experimental association rate constant (k_on_) and involved a massive amount of computation (2 ms of aggregate simulation time on the GPUgrid computing network), the Markov state model necessitates the use of a significant lag time (in this study, 110 ns), which fundamentally limits the ability to resolve features of the binding mechanism below such timescales (*e.g.* short-timescale protein and water dynamics). Barnase-barstar association has also been studied using Brownian dynamics simulations with rigid, atomistic protein models and implicit solvent.^7, 8^

Here, we applied the weighted ensemble (WE) path sampling strategy^9^ to orchestrate MD simulation, yielding atomically detailed pathways and rate constants for the barnase-barstar binding process with <1% of the aggregate simulation time that was generated in the Markov state model study mentioned above.^6^ A hallmark of the WE strategy is that pathways and rate constants can be generated for rare events (*e.g.*, protein folding and binding) using any type of stochastic dynamics^10^ in orders of magnitude less computing time than standard simulations.^11–16^ Furthermore, this efficiency increases exponentially with the effective free energy barrier of the rare event.^11^ Importantly, the generated pathways are fully continuous, ensuring that all sampled states of the rare-event process are “on pathway.” The WE strategy has enabled atomistic simulations of a protein-peptide binding process with implicit solvent^1^ as well as the estimation of the slow “basal” k_on_ (beyond the tens of milliseconds timescale) for barnase-barstar association in the absence of electrostatic interactions^13^ using flexible, residue-level models.^17^ The present study is the most ambitious WE application to date, yielding a diverse ensemble of atomically detailed protein-protein binding pathways in explicit solvent.

The barnase-barstar system has been a classic system for studying protein-protein binding processes due to the rapid, diffusion-controlled k_on_,^18^ The two proteins are produced by the bacteria, *Bacillus amyloliquefaciens*, with barstar being the “safety pin” that prevents barnase from unleashing its lethal “grenade” of ribonuclease activity inside the cell.^19^ The rapid k_on_ has been attributed to long-range electrostatic interactions between the negatively charged barnase-binding surface of barstar with the positively charged active site of barnase.^18^ Furthermore, barstar has been found by a theoretical study to be electrostatically optimized for tight binding to barnase in terms of the balance between protein interactions and desolvation.^20^

## 2. Methods

### 2.1 Weighted Ensemble (WE) Simulations

#### Overview of the WE strategy

To enable direct atomistic simulations of the protein-protein binding process, we applied the weighted ensemble (WE) path sampling strategy^9^ in conjunction with MD simulations. This strategy involves carrying out a large number of trajectories in parallel, with each trajectory assigned a statistical weight to properly represent the path ensemble. To control the distribution of trajectories, configurational space is divided into bins along a progress coordinate towards the target state and trajectories are evaluated at fixed time intervals *τ* for resampling. The resampling procedure involves either the replication or pruning of trajectories to maintain a specified number of target trajectories/bin while adjusting trajectory weights according to rigorous statistical rules such that no bias is introduced into the dynamics. The WE strategy can be applied under steady-state or equilibrium conditions. The former is maintained by recycling trajectories that reach the target state by terminating such trajectories and starting new trajectories from the initial state with the same trajectory weights. In the present study, the WE strategy was applied under equilibrium conditions with no recycling of trajectories to permit the refinement of key states after completion of the simulation for calculations of rate constants.^21^ All WE simulations were carried out using the open-source, highly scalable WESTPA software (https://westpa.github.io/westpa).^22^

#### General simulation strategy

To generate a diverse ensemble of binding pathways, our simulation strategy featured the following: (i) providing multiple chances for each of 100 pre-equilibrated unbound states to result in successful binding pathways by initiating 16 trajectories from each unbound state to yield a total of 1600 trajectories for the binding simulation, and (ii) “front-loading” the simulation among bins between the unbound state and metastable “encounter complex” intermediate by maintaining a constant number of 1600 trajectories across all bins of the progress coordinate at a given WE iteration thereby reducing the likelihood of trajectories from a particular unbound state to be pruned early in the simulation.

#### Generation and pre-equilibration of unbound states

Prior to carrying out WE simulations of the binding process, representative unbound conformations of each binding partner were generated by running a separate WE simulation starting from the conformation of that partner in the native, bound complex. Each “preparatory” WE simulation was carried out under equilibrium conditions, which refers to no recycling of trajectories as mentioned above, using a one-dimensional progress coordinate that consisted of the heavy-atom RMSD of the protein from its conformation in the crystal structure of the protein-protein complex. This RMSD coordinate was divided into 45 bins, with finely spaced bins every 0.1 Å from 0 to 3.0 Å, more coarsely spaced bins every 0.5 Å from 3.0 to 10 Å, and a single bin for all values ≥10 Å. The simulations were carried out using a target number of 12 trajectories/bin for 1200 WE iterations with each iteration having a fixed length *τ* of 5 ps, yielding a maximum trajectory length of 6 ns to achieve steady probability distributions as a function of the progress coordinate. Unbound states for the binding simulations were then generated by selecting conformations of each binding partner according to its probability from the last iteration of the preparatory WE simulation and randomly orienting the partners with respect to each other at a minimum separation of 20 Å to yield 1728 pairs of unbound conformations of barnase and barstar. These 1728 pairs were then reduced to 100 pairs by assigning trajectories to appropriate bins along the minimum separation distance dimension of the two-dimensional binding progress coordinate (see below) and combining the lowest-weight trajectories according to the standard WE algorithm. Each of the resulting 100 unbound states was then re-immersed in dodecahedral boxes of water molecules, equilibrating the solvent in two stages as described below under “propagation of dynamics.”

#### Binding simulation

To simulate the binding process, a WE simulation was carried out under equilibrium conditions by initiating 16 separate trajectories from each of the 100 pre-equilibrated unbound states with appropriately renormalized weights for all 1600 trajectories. A two-dimensional progress coordinate was used throughout the binding simulation, consisting of (i) the minimum separation distance between and two binding partners, and (ii) the heavy-atom RMSD of the two barstar “anchor” residues relative to the barnase-bound crystal pose following alignment on barnase. We define the anchor residues – D35 and D39 – as the residues of barstar that become the most buried upon binding barnase. The first dimension of the progress coordinate was divided into two bins to separate conformations with the binding partners in van der Waals contact (<5 Å) and not in contact (≥5 Å). Prior to collisions to form the encounter complex, the second dimension of the progress coordinate was divided into a total of 72 bins with coarsely spaced bins every 1 Å from 10 to 60 Å and finely spaced bins every 0.5 Å from 0 to 10 Å. To make optimal use of a given number of CPU cores, the total number of trajectory segments that were being carried out at a time was fixed at a constant number of 1600, adjusting the number of target trajectories in each bin as appropriate. After forming the encounter complex, these adjustments resulted in an average of 22 target trajectories/bin. To obtain a steady value of the k_on_ (Fig. S1 Supporting Information), the binding simulation was carried out for 650 WE iterations with a fixed interval *τ* of 20 ps for each iteration, yielding a maximum trajectory length of 13 ns and an aggregate simulation time of 18 μs. All analysis was performed every *τ* unless otherwise noted.

#### Propagation of dynamics

Dynamics of the WE simulations were propagated using the Gromacs 4.6.7 dynamics engine^23^ with the Amber ff03* force field^24^ and TIP3P water model.^25^ Heavy-atom coordinates for initial models of the unbound proteins and native complex were taken from the crystal structure of the barnase-barstar complex (PDB code: 1BRS).^26^ Hydrogen atoms were added to each model using ionization states present in solution at pH 7. Each system was immersed in a sufficiently large dodecahedron box of explicit water molecules to provide a minimum 12 Å clearance between the solutes and box walls for the unbound states in which the binding partners were separated by 20 Å. A total of 31 Na+ and 29 Cl-ions were included to neutralize the net charge of the protein system and to yield the experimental ionic strength (50 mM).^18^ The entire simulation system consisted of ~100,000 atoms.

Prior to carrying out WE simulations, the systems were first subjected to energy minimization and then two stages of equilibrating the solvent while applying harmonic constraints to the proteins with a force constant of 10 kcal mol^−1^ • Å^−2^. During the first stage, the system was equilibrated for 20 ps at constant temperature (25 °C) and volume. During the second stage, the system was equilibrated for 1 ns at constant temperature (25 °C) and pressure (1 atm). Since the WE strategy requires stochastic dynamics,^10^ a stochastic thermostat was used, *i.e.* velocity rescaling thermostat^27^ with a coupling constant of 0.1 ps; pressure was maintained using a weak Berendsen barostat^28^ with a coupling constant of 0.5 ps. To enable a 2-fs time step. bonds involving hydrogens were constrained using the LINCS algorithm.^29^ Van der Waals interactions were switched off smoothly between 8 and 9 Å along with the application of a long-range analytical dispersion correction to energy and pressure. Real-space electrostatic interactions were truncated at 10 Å and long-range electrostatic interactions were calculated using particle mesh Ewald summation.^30^

### 2.2 Calculation of rate constants and percentage of productive collisions

To calculate rate constants from the binding simulation, the resulting equilibrium set of trajectories can be decomposed into two steady states, “AB” and “BA”.^21^ The “AB” steady state consists of trajectories in the binding direction most recently in state A (*e.g.* unbound state) than in B (*e.g.* encounter complex), and the “BA” steady state consists of trajectories in the unbinding direction most recently in B than in A. Bimolecular and unimolecular rate constants *k_ij_* between states *i* and *j* of the binding process were calculated using the following:^21^

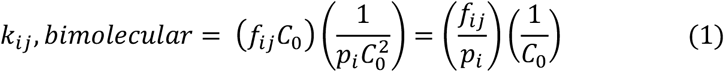

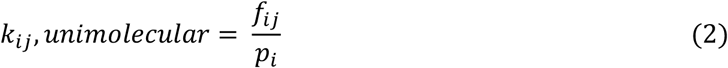

where *f_ij_* is the flux of probability carried by trajectories originating at any point in the simulation in state *i* and arriving in state *j*, *p*_*i*_ is the fraction of trajectories more recently in state *i* than in *j*, and *C*_0_ is the reference concentration of the binding partners (1.7 mM), calculated as 1/(*N_A_V*) where *N_A_* is Avogadro’s number and *V* is the volume of the dodehedral simulation box (956 Å^3^).

Equation (1) was used for computing the bimolecular k_on_ as well as the rate constant for formation of the encounter complex (k_1_). Equation (2) was used for computing the unimolecular rate constant for rearrangement of the encounter complex to the bound state (k_2_). All uncertainties in rate constants represent 95% confidence intervals and were estimated using a Monte Carlo blocked bootstrapping technique.^9, 31^ The rate constant k_1_ was calculated using the entire simulation while k_2_ and k_on_ were calculated using the latter half of the simulation during which greater sampling was focused on the rearrangement of the encounter complex to the bound state.

The percentage of productive collisions (*i.e.*, encounter complexes that succeed in rearranging to the bound state) was calculated, as done before,^32^ according to the following equation:

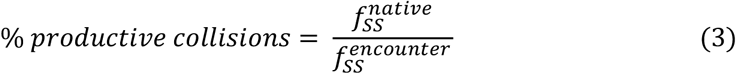

where 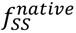 is the steady-state flux into the native, bound state and 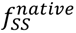 encounter is the steady-state flux into the encounter complex; both fluxes were evaluated only after an approximate steady state was achieved. The uncertainty in the % productive collisions represents a 95% confidence interval, which was estimated by propagation of error in the steady state fluxes; the latter was calculated using the Monte Carlo block bootstrapping technique mentioned above.

#### State definitions

State definitions used for calculations of rate constants were determined from the probability distribution of the binding simulation as a function of the progress coordinate yielded by WE itself (Fig. 1A). The unbound state was defined as having a minimum separation distance of ≥ 20Å between the proteins. The encounter complex intermediate was defined to include only non-native complexes that had a sufficiently long survival time to proceed to the native complex, *i.e.* heavy-atom RMSD of ≥ 4 Å and ≤ 20 Å for D35 and D39 of barstar after alignment on barnase and a minimum separation distance of ≤ 3 Å between the two proteins. The bound state was defined as having a heavy-atom RMSD ≤ 3.5 Å for D35 and D39 of barstar after alignment on barnase and a minimum separation distance of ≤ 3 Å between the two proteins.

**Fig. 1.**
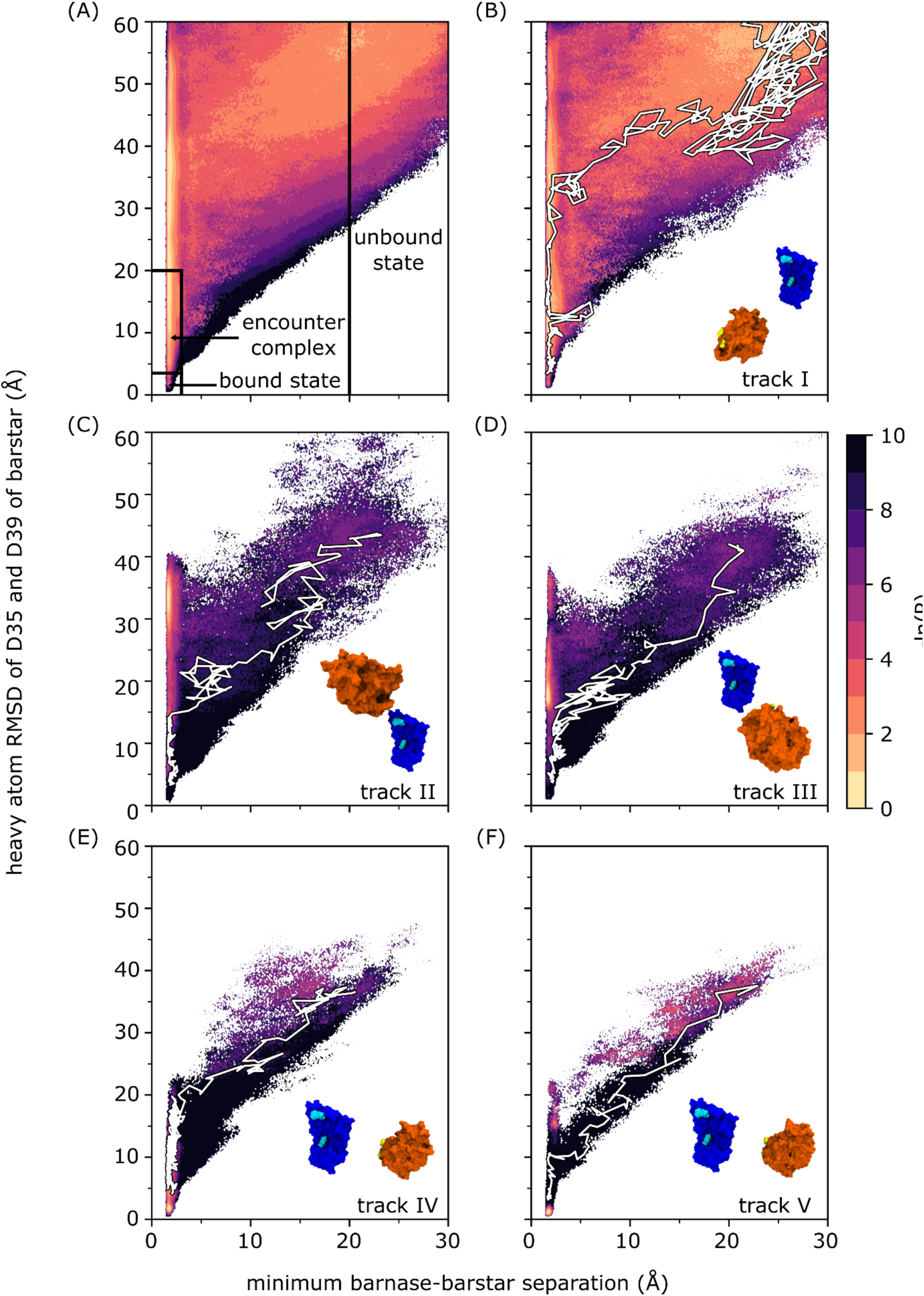
Probability distributions as a function of the WE progress coordinate for **(A)** all simulation data with definitions of the unbound state, encounter complex, and bound state delineated by solid black lines, and **(B-E)** tracks I-V. In each of the panels B-E, a snapshot of the initial unbound states is included with molecular surface representations of barnase and barstar in blue and orange, respectively, barnase anchor residues (K27 and R59) are highlighted in cyan, and barstar anchor residues (D35 and D39) are highlighted in yellow; a representative binding pathway along the corresponding probability distribution is highlighted in white. A movie of the representative pathway in Fig. 1B is included in the Supporting Information (Movie S1). The probability distribution of each binding track was normalized according to the total probability of that track. The color scale represents -lnP where P is the probability density calculated as the sum of appropriate trajectory weights.

### 2.3 Maps of ligand entry point distributions and conformation space networks

Inspired by Dickson and Lotz,^33^ we generated spherical maps of ligand entry points and constructed conformation space networks to visualize the evolution of various properties along the ensemble of simulated binding pathways.

Spherical maps were constructed by projecting the probability distribution of ligand entry points for diffusional collisions of the barstar ligand and barnase receptor onto a unit sphere that is centered on the barnase receptor. The receptor was rotated such that W44 of barnase is aligned with the z-axis and R59 of barnase is aligned with the y-axis. The probability distribution was generated by creating a histogram using 30 bins and trajectory weights at each ligand entry point.

Conformation space networks were constructed by first applying the KCenters clustering algorithm to conformations of the two longest pathways for each of the 203 binding events to yield a total of 2000 clusters. The clustering was carried out using a Canberra distance metric as implemented in the MSMBuilder software package^34^ and a feature vector that consisted of the WE progress coordinate used for the binding simulation. Conformational space networks were then generated using the Gephi 0.9.2 software package^35^ and ForceAtlas 2 layout algorithm,^36^ with each node represent a cluster center and the edges between nodes representing observed transitions between each cluster. The size of each node is proportional to the total statistical weight over all conformations in the corresponding cluster and colored according to the weighted average of the property of interest over all conformations of that cluster. The committor probability for each cluster was calculated from the number of transitions between relevant nodes.

### 2.4 Calculation of pairwise residue contact maps

To identify kinetically important residues for the binding process, we searched for the most probable intermolecular residue contacts in the transition path ensemble (TPE), which consists of only the transient portions of the productive binding pathways between the stable states. These so-called “transition paths” begin where the trajectory last exits the initial unbound state and end where the trajectory first enters the bound state. A pair of residues was considered to be in contact if any of their heavy atoms were ≤ 4.5 Å of each other. The probability of forming a pairwise residue contact in the TPE was calculated by summing over the statistical weights of all TPE conformations where the two residues are in contact; the weight of each TPE conformation was calculated by summing over the weights of its successful child trajectories.

### 2.5 Calculation of sidechain conformational entropy per residue

The sidechain conformational entropy *S*_*X*_ of each residue in a given state *X*(*i.e.* unbound state, encounter complex, or bound state) was calculated using the following:

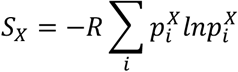

where *R* is the ideal gas constant and 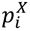 is the probability of observing a particular set *i* of *χ* angles for the sidechain of the residue in state *X* in the WE simulation; 25 bins were used to generate the histogram of each probability distribution.

### 2.6 Monitoring protein desolvation and percent burial of individual residues

To monitor protein desolvation during the binding process, we tracked the number of water molecules *N*_*w*_ within 6 Å of each protein to encompass the first two solvation shells. We then calculated a “percent solvation” by dividing the average *N*_*w*_ in a particular conformation by the average *N*_*w*_ in the unbound state.

Percent burials of barstar residues, D35, D39, W38, and W44, upon binding were calculated as (SASA in the selected configuration)/(average SASA in the unbound state) x 100% where the SASA is the solvent accessible surface area and calculated using the Shrake and Rupley algorithm^37^ as implemented in MDTraj Python library.^38^

### 2.7 Calculation of percent occupancies of interfacial crystallographic water positions

The percent occupancy of each of the nine interfacial, crystallographic water positions was calculated for the bound-state ensemble by summing over the probabilities of all trajectories in which a water molecule forms occupies one of the nine positions, forming hydrogen bonds with the corresponding residues in each of the two proteins. Trajectory probabilities were normalized by the population of the bound-state ensemble and included in the sum of probabilities if a given position is occupied at any point within each WE iteration of fixed length *τ* = 20 ps. Hydrogen bond formation was monitored every ps and defined as having a donor-acceptor distance of ≤ 3 Å and a donor-acceptor angle of ≥ 90º using the MDAnalysis Python library.^39^

## 3. Results & Discussion

### 3.1 Mechanism of binding

Our weighted ensemble (WE) simulation was successful in generating a large ensemble of fully continuous, atomically detailed protein-protein binding pathways in explicit solvent. In particular, a total of 203 binding pathways involving the barnase and barstar proteins were captured by a single WE simulation within 30 days using 1600 CPU cores at a time on XSEDE’s Stampede supercomputer. Notably, the aggregate simulation time was 18 μs, which is <1% of that generated in a prior atomistic simulation study of barnase-barstar binding using a Markov state model.^6^

Results reveal a two-step binding process in which a nonnative “encounter complex” (Fig. 1A) is first formed, followed by rearrangement of this metastable intermediate to the native, bound state. While only 4% of the aggregate simulation time yielded successful binding pathways, 81% resulted in diffusional collisions and 35% resulted in the formation of encounter complexes, which are defined here as productive end points of diffusional collisions that eventually rearrange to the bound state. Among all of the collisions, only 11 ± 5% were productive. We note that all percentages reported in this study represent WE weighting.

As shown in Table 1, our computed association rate constant k_on_ [(2.3 ± 1.0) x 10^8^ M^−1^s^−1^] is within error of experiment [(2.86 ± 0.7) x 10^8^ M^−1^s^−1^].^18^ Given that the computed rate constant for formation of the encounter complex k_1_ [(1.8 ± 0.2) x 10^9^ M^−1^s^−1^] is approximately equal to the k_on_ and that the computed rate constant k_2_ for rearrangement of the encounter complex to the bound state is relatively fast [(2.7 ± 0.5) x 10^9^ s^−1^], the rate-limiting step for the binding process is the diffusion-controlled formation of the encounter complex. The rate constant k_1_ for this initial step is on the order of the Smoluchowski limit (~5 x 10^9^ M^−1^s^−1^) despite the orientational constraints due to electrostatic interactions between the proteins.^40^ Since our WE simulation was set up to focus the sampling primarily on binding events, we did not observe a sufficient number of unbinding events to compute statistically robust rate constants in the unbinding direction and therefore focused exclusively on kinetics in the binding direction.

**Table 1:** Computed rate constants and 95% confidence intervals for the barnase-barstar binding process.

| process | rate constant | value |
| --- | --- | --- |
| unbound state → encounter complex | $k_1 (\text{M}^{-1}\text{s}^{-1})$ | $1.8 \pm 0.2 \times 10^9$ |
| encounter complex → bound state | $k_2 (\text{s}^{-1})$ | $2.7 \pm 0.5 \times 10^{10}$ |
| unbound state → bound state | $k_{\text{on}} (\text{M}^{-1}\text{s}^{-1})$ | $2.3 \pm 1.0 \times 10^8$ |
| | experimental $k_{\text{on}} (\text{M}^{-1}\text{s}^{-1})$<sup>18</sup> | $2.86 \pm 0.67 \times 10^8$ |

### 3.2 Diversity of binding pathways

Our simulation generated a diverse ensemble of binding pathways among five separate “tracks” that each originated from different configurations of the unbound state and are therefore independent with no common trajectory segments (Fig. 1B-F). These tracks, referred to as tracks I to V, vary in the extent to which the binding partners must rotate relative to each other in order to collide and form productive encounter complexes. Track I (Fig. 1B) is the most indirect binding track, requiring the greatest amount of relative rotations, whereas tracks IV and V are the most direct binding tracks, requiring the least amount of relative rotations. Despite the fact that the unbound configurations for tracks IV and V are very similar, track V yielded less direct pathways than track IV to forming the encounter complex.

The most indirect track (track I, Fig. 1B) is the most probable track, accounting for the lion’s share of the probability distribution that was sampled by the simulation as a function of the WE coordinate (Fig. 1A). Furthermore, the distribution of event duration times, or times required to traverse transition paths, is essentially identical for the most indirect track and the full set of binding pathways with the most probable event duration being 6.7 ns (Fig. S2 in the Supporting Information). The fact that the most indirect binding pathways are the most probable pathways in our simulation is not surprising given that the binding interfaces of partner proteins are not highly likely to be pointing directly at each other in their unbound states and must therefore involve rotations of the proteins in order to productively collide to form encounter complexes that eventually rearrange to the native complex. It is worth noting that the short-timescale protein and water dynamics of all binding pathways captured by our simulation would not be resolved in analysis of Markov state models given the long lag times required for these models (*e.g*., 110 ns for previous simulations of barnase-barstar binding).^6^

Given that only subtle differences exist between high-resolution crystal structures of the barnase-barstar complex^26^ and those of the corresponding unbound proteins,^41, 42^ it might appear that the complex forms by rigid-body association. However, our simulations reveal a much more dynamic view of the structures as the two proteins perform their molecular dance towards a final “embrace” to form the native complex. As evident in a representative movie of the most indirect track (track I) and more importantly, the most probable track, the sidechains are highly dynamic in both proteins even in the bound-state ensemble (Movie S1 in the Supporting Information).

### 3.3 Kinetically important interactions

To identify kinetically important interactions for the binding process, we constructed a map of probabilities for forming each possible pair of intermolecular residue contacts in the transient states that comprise the transition path ensemble (TPE; see Methods). As shown in Fig. 2A and 2B, the most probable contacts are D35bs-R59bn and W44bs-S38bs (“bn” for barnase and “bs” for barstar), which are formed 34% and 23% of the time, respectively. These two contacts fasten both edges of the barnase binding interface in the encounter complex such that the subsequent rearrangement of the encounter complex to the bound state reduces the conformational entropy of R59bn more than any other residue (Fig. 2C).

**Fig. 2.**
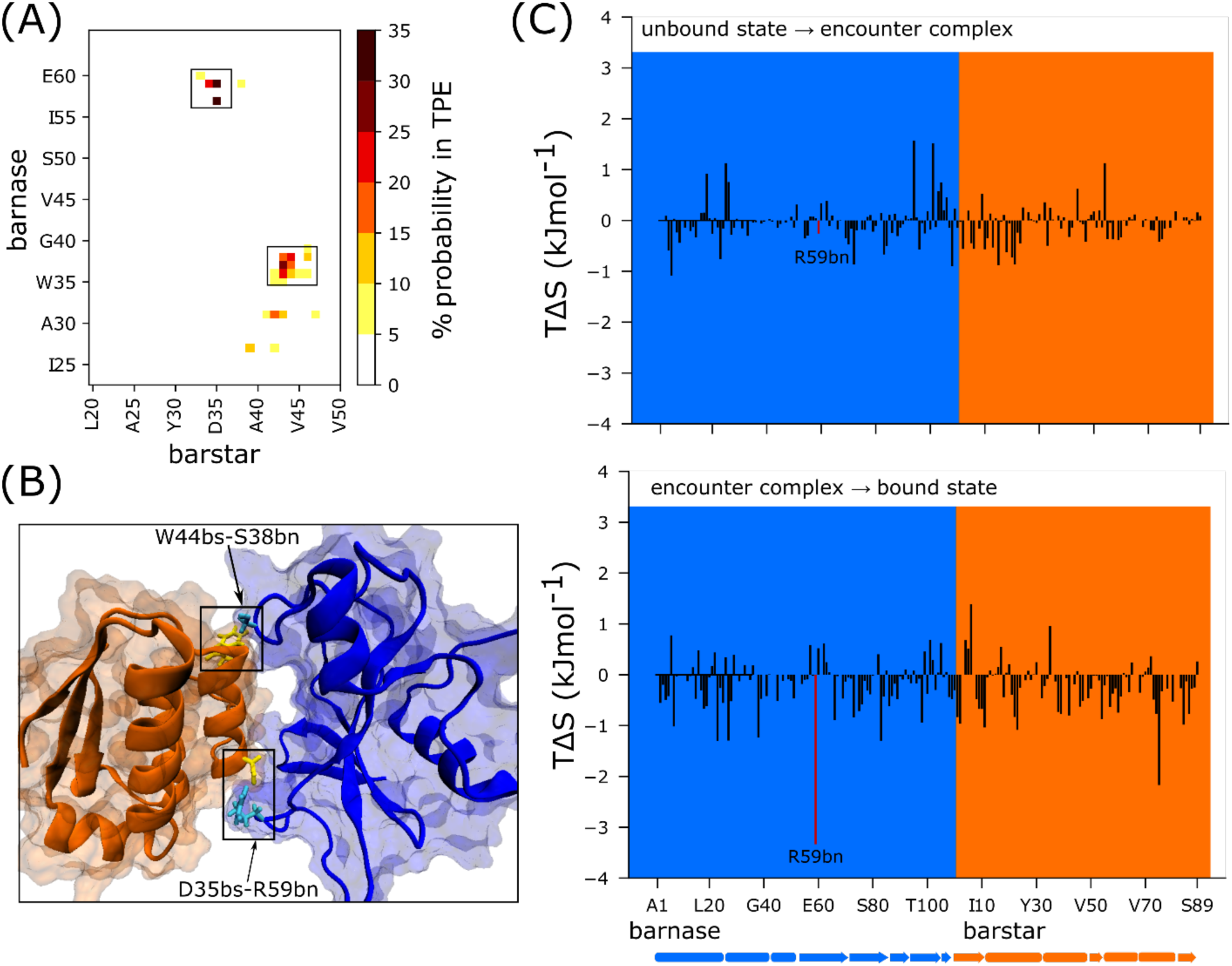
**(A)** Map of pairwise residue contacts formed in the encounter complex ensemble. The color bar represents the percent probability of forming a given pair of residue contacts in the transition path ensemble (TPE). **(B)** Locations of the most kinetically important residue contacts indicated in the crystal structure of the native complex of barnase (blue) and barstar (orange) with R59bn and S38bn in cyan and D35bs and W44bs in yellow. **(C)** TΔS per residue where ΔS is the change in sidechain conformational entropy of the residue upon forming the encounter complex (top) and upon rearrangement of the encounter complex to the bound state (bottom). Barnase and barstar residues are indicated by the blue and orange shaded regions, respectively. Highlighted in red is R59bn, the residue with the largest TΔS upon rearrangement of the encounter complex to the native complex.

The importance of R59bn for the kinetics of binding is also evident from an analysis of the encounter complex ensemble. In particular, among the diverse relative orientations of the binding partners that resulted in collisions to form encounter complexes, productive collisions generally involved contacts with R59bn or other residues in its vicinity (Fig. 3). As shown in Fig. 4, a representative pathway along the most indirect binding track, track V, involves formation of the D35bs-R59bn contact upon collision of barnase and barstar followed by formation of the W44bs-S38bn contact in the encounter complex thereby fastening both edges of the barnase binding interface before rearranging to the native complex. Our results reveal that the barstar residues, D35bs and D39bs, are not only the most buried upon binding barnase, but become buried earlier than other barstar residues with D35bs burying earlier than D39bs (Fig. S3A and S3B). In addition, both Trp residues at the barnase binding interface, W38bn and W44bn, become buried upon forming the encounter complex with W44bn becoming buried before W38bn (Fig. S3C and S3D). Thus, the detection of a change in Trp fluorescence by stopped-flow experiments would result from formation of both the encounter complex and native complex.

**Fig. 3.**
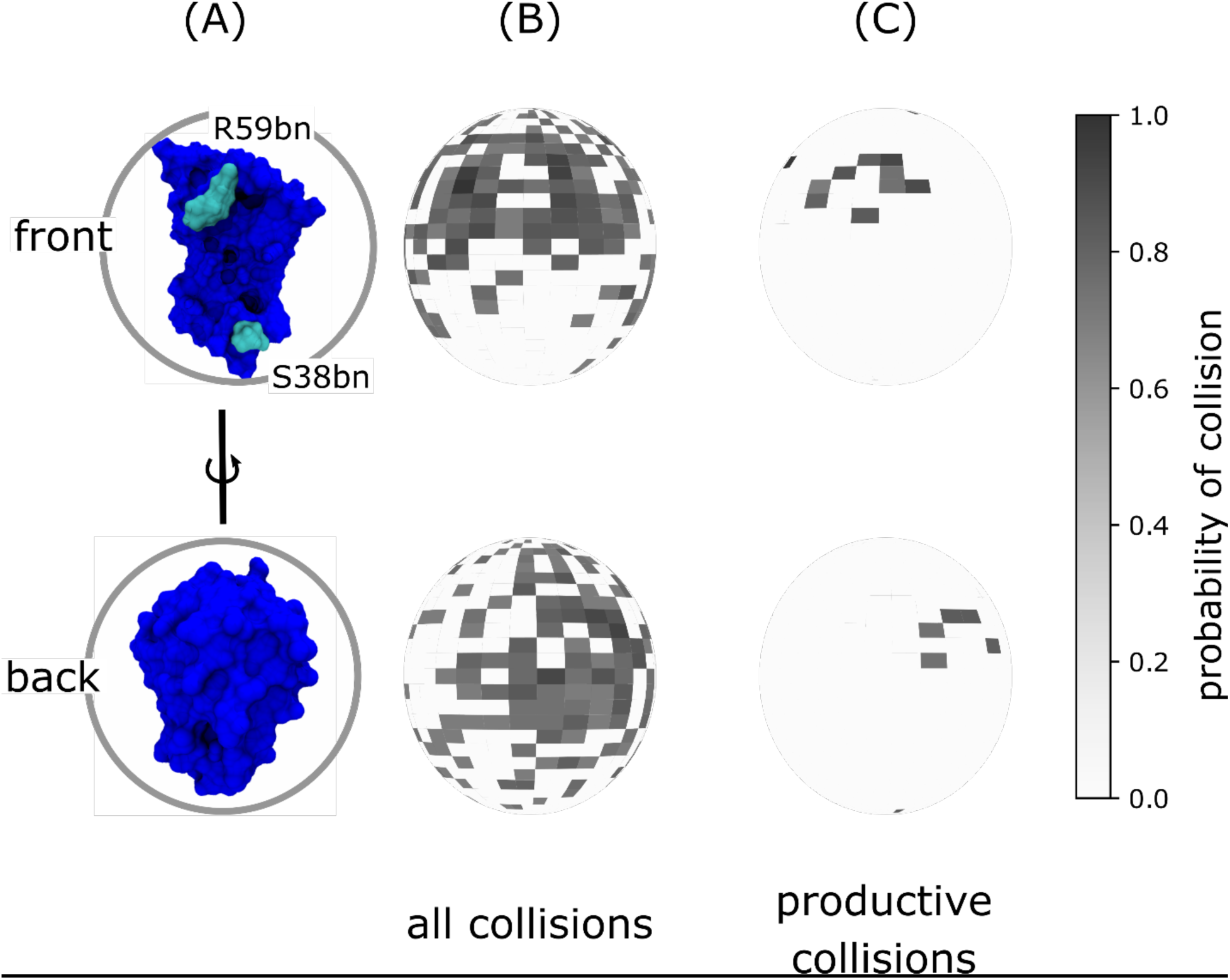
Spherical maps of ligand entry point distributions for the barstar ligand in diffusional collisions with the barnase receptor. **(A)** Reference orientations of barnase (blue) for ligand entry point distributions in panels B and C where each probability distribution is projected onto a unit sphere that is centered on barnase in one of two orientations: the front view corresponding to a “head-on” view of the barnase binding interface and the back view corresponding to a 180° rotation around the vertical axis of the front view. **(B)** Ligand entry point distributions for all diffusional collisions. **(C)** Ligand entry point distributions for only productive collisions that form encounter complexes, which subsequently rearrange to the bound state. As shown in panel C, the most probable entry points are in the vicinity of R59bn (cyan in panel A), which lies at one edge of the barnase binding interface, as opposed to S38bn, which lies at the other edge.

**Fig. 4.**
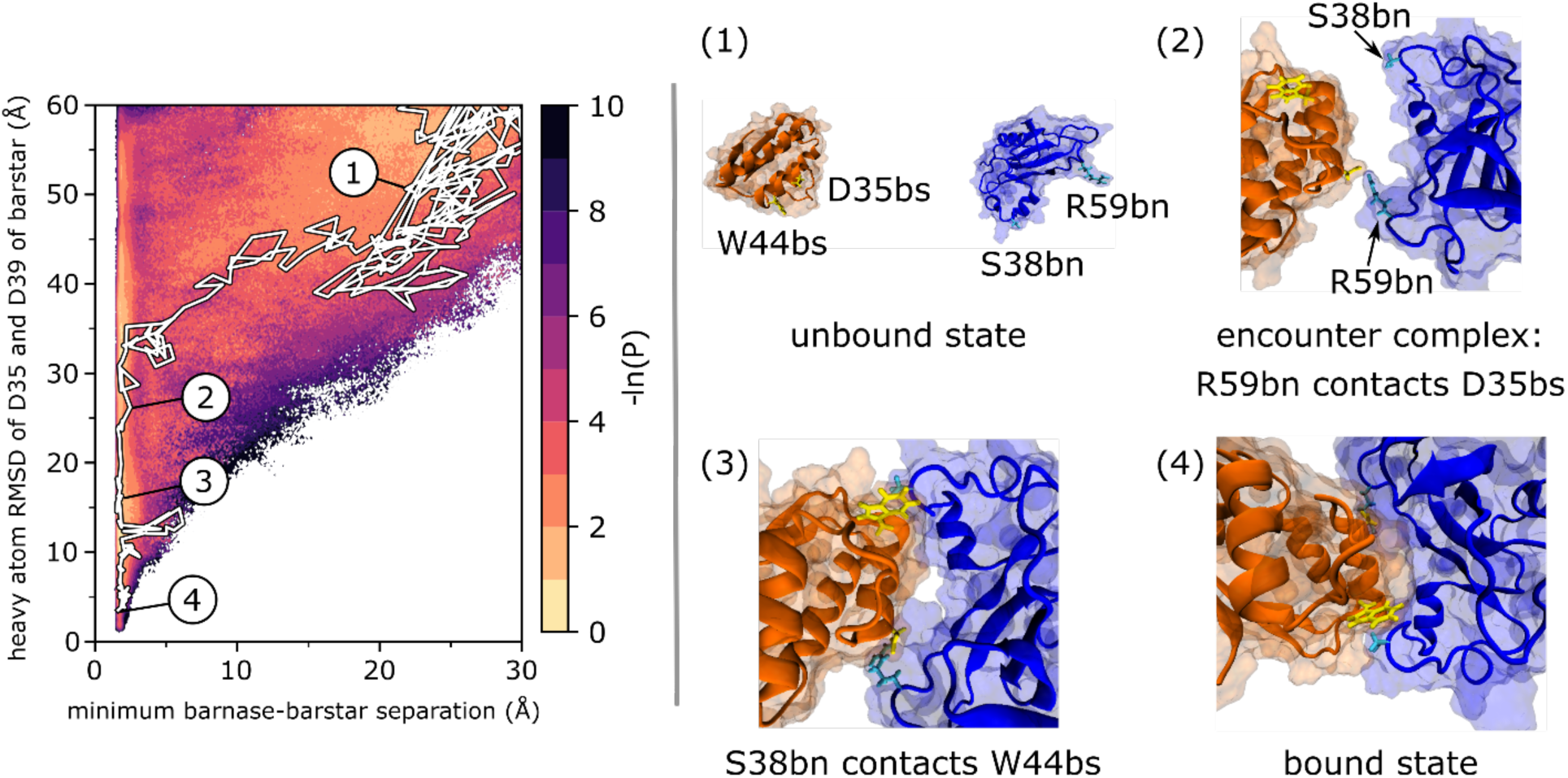
A representative, continuous binding pathway of track I superimposed onto the probability distribution over the WE progress coordinate. The simulation was initiated from an unbound state in which the binding partners were separated by 20 Å (1). Diffusional collision to form an encounter complex that includes forming initial native D35bs-R59bn contacts (2), followed by securing native W44bs-S38bn contacts during rearrangement of the encounter complex (3) to the native, bound state (4). This trajectory is also illustrated in Movie S1 with conformations sampled every ps.

Our identification of R59bnand S38bn as kinetically important residues for the barnase-barstar binding process is consistent with previous studies. In particular, experimental studies have identified R59bn as the most kinetically important barnase residue for the binding process of barnase and barstar, resulting in a >9-fold slower k_on_ when mutated to an alanine.^43^ Furthermore, S38bn was found to form intermolecular contacts in the majority of encounter complexes sampled by a recent simulation study that involved a different force field (Amber ff99SB-ILDN^44^ and TIP3P^25^) and construction of Markov state models.^6^

### 3.4 Evolution of solvent configuration during the binding process

To determine when the two proteins undergo desolvation of their binding interfaces before forming the native complexes, we monitored the percent solvation of each conformation relative to the unbound state, tracking the number of water molecules within 6 Å of each protein (see Methods). We then generated a conformational space network to visualize the various binding tracks and colored this network according to the minimum percent solvation thereby detecting *any* instance of desolvation. As shown in Fig. 5B, protein desolvation occurs in the late stages of the binding process in our simulations. In particular, the two proteins undergo the greatest extent of interface desolvation during the rearrangement of the encounter complex to the native complex. This result is consistent with an experimental study in which the characterization of the transition state for the rearrangement of the encounter complex to the native complex revealed that most of the interface desolvation has not yet occurred based on a low activation entropy.^45^

**Fig. 5.**
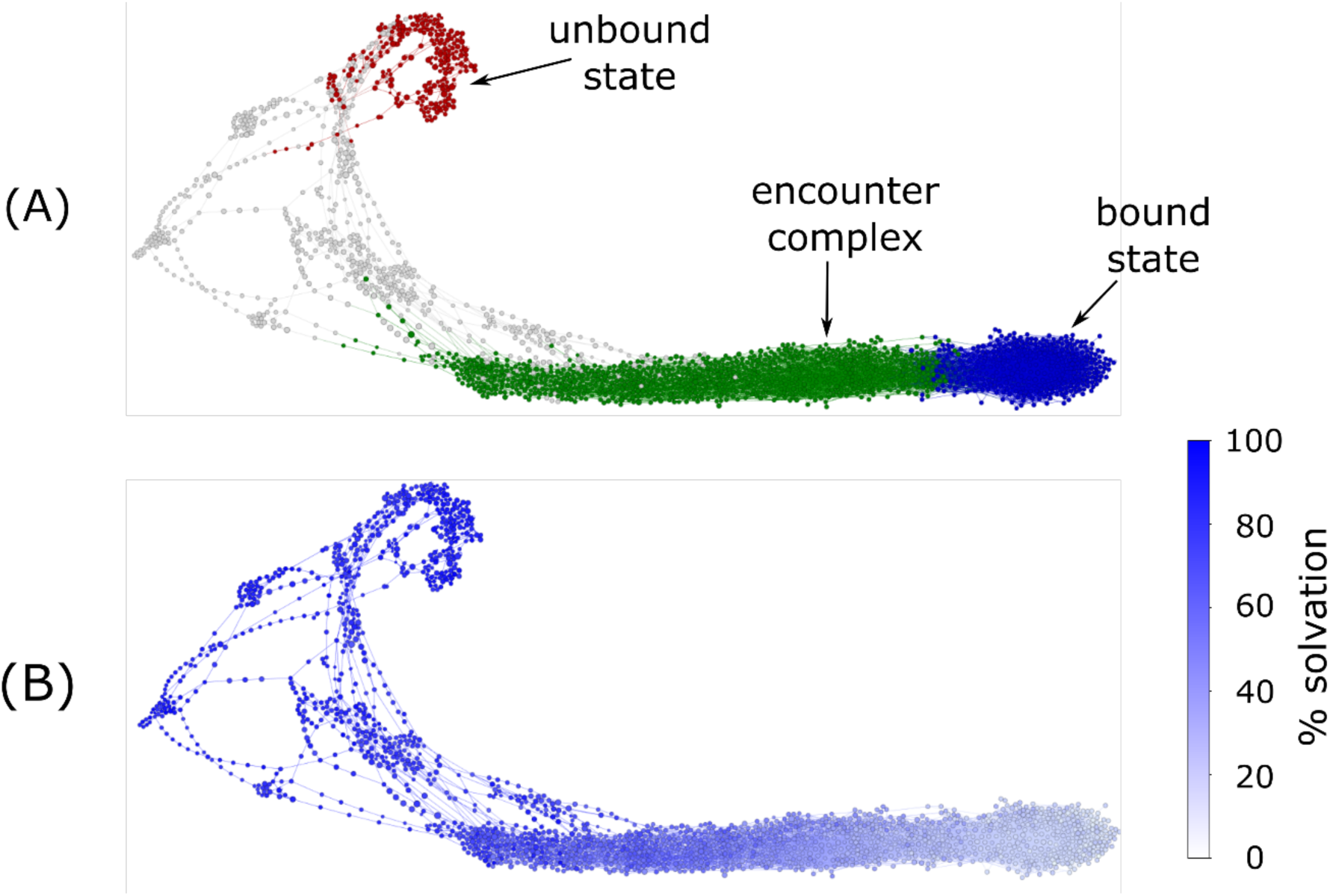
Conformational space networks of barnase-barstar binding pathways colored according to **(A)** state definitions used for calculations of rate constants, and **(B)** percent solvation in a given cluster of conformations (see Methods).

Notably, our simulations revealed that during the binding process, water molecules from the bulk solvent eventually occupy the positions of all but one of the nine interfacial crystallographic water molecules that bridge hydrogen bonds between barnase and barstar in the native complex.^26^ As shown in Table 2, the eight positions are occupied 4% to 48% of the time. The occupancy of these positions is an encouraging validation of the force field and water model – particularly since the simulations were started from the unbound state. Furthermore, these results suggest that the interfacial water molecules in the crystal structure of the native complex are present in solution as well as the crystal environment. Finally, our simulation identified water molecules in the bound state ensemble that are not resolved in the crystal structure of the native complex and bridge hydrogen bonds between residues that we have identified as kinetically important (Fig. S4).

**Table 2.** Percent occupancies of solvent in the simulated bound-state ensemble at positions of crystallographic waters that bridge hydrogen bonds between residues in barnase (bn) and barstar (bs). Crystallographic waters are listed in order of top to bottom along the corresponding positions (red spheres) at the binding interfaces of barnase (blue) and barstar (orange).

| *Solvent* | *Solvent ASA (Å<sup>2</sup>)* | *Residues bridged by hydrogen bonds of the water molecule* | *% occupancy in the simulated bound-state ensemble* |
| --- | --- | --- | --- |
| *Wat128* | 0 | K62bn/Y103bn – D35bs | 17 |
| *Wat22* | 0 | K62bn/N58bn – D35bs | 22 |
| *Wat29* | 0 | R59bn – D35bs | 21 |
| *Wat14* | 0 | E73bn – D35bs | 0 |
| *Wat48* | 3 | I55bn/E73bn – W38bs | 33 |
| *Wat33* | 0 | K27bn/E73bn – D39bs | 48 |
| *Wat155* | 1 | R83bn – G43bs | 6 |
| *Wat36* | 9 | S38 – V45bs | 11 |
| *Wat93* | 17 | S38bn – Y47bs | 4 |

### 3.5 Choice of progress coordinate for protein binding processes

As described in Methods, our WE simulation of the protein-protein binding process employed a two-dimensional progress coordinate consisting of (i) the minimum separation distance between barnase and barstar, and (ii) the heavy-atom RMSD of the two barstar “anchor” residues, D35 and D39, relative to the barnase-bound crystal pose following alignment on barnase. The first dimension was essential for avoiding the merging of trajectories in which the binding partners are in contact with trajectories in which the binding partners are not in contact thereby ensuring sufficient sampling of the encounter complex and enabling converged estimates of rate constants k_1_ and k_2_ as well as the percentage of productive collisions. The second dimension was intended to generate all types of collisions (productive and nonproductive) starting from a diverse set of relative orientations of the binding partners in their unbound states and to distinguish between the nonnative encounter complex from the native bound state.

Although the RMSD of the entire native complex has been previously found to be a poor descriptor for monitoring the progress of protein-protein association pathways,^7^ we have demonstrated that the RMSD can be an effective progress coordinate if the deviations are calculated for only the ligand after alignment of the protein receptor. At large ligand-receptor separations, this “binding” RMSD functions as a distance metric and at shorter separations, as a metric of both distance and relative orientations of the binding partners with respect to those of the bound state. For both this study and our recent atomistic WE study of protein-peptide binding,^1^ we have focused on taking the RMSD of the ligand “anchor” residues, which are the residues that become the most buried upon binding the protein receptor. Such anchor residues have been proposed to smooth out the binding process by avoiding kinetically costly structural rearrangements.^46^ We note that the main limitation of WE and related strategies (see review^47^) is that the free energy barriers to be surmounted may be orthogonal to the selected progress coordinate. In principle, however, if the progress coordinate captures the slowest relevant motion, then faster, correlated coordinates will also be captured.^48^

A future goal for the development of WE strategies is to generate a greater diversity of indirect binding pathways, including those that might involve rearrangement of the encounter complex to the bound state via “crawling” of one protein over the molecular surface of the partner protein. Promising strategies for achieving this goal are the use of progress coordinates that more extensively cover configurational space (*e.g.* adaptive construction of bins using Voronoi procedures^10^) and the improvement of schemes for replication and pruning of trajectories to maximize the number of distinct unbound states that yield successful binding pathways.

## 4. Conclusions

In closing, we have demonstrated the power of the weighted ensemble (WE) strategy^9^ in enabling explicit-solvent MD simulation of a protein-protein binding process. Our simulation involves the prototypical barnase-barstar system for studying protein-protein binding and provides several insights regarding the binding mechanism that cannot be obtained by laboratory experiments.

First, our simulation provides atomically detailed views of the binding pathways, including states that are too transient to be captured by experiment. In total, 203 binding pathways were generated, yielding a computed k_on_ that is in good agreement with experiment.^18^ Among the diverse path ensemble, the most probable binding pathways were the most indirect, requiring the greatest extent of rotation of the partner proteins in order to collide productively. Our simulations reveal that both proteins are much more dynamic than expected from the subtle differences between the high-resolution, experimental structures of the corresponding unbound and partner-bound states^26, 41, 42^ with highly dynamic sidechains in both proteins throughout the binding process.

Second, our simulation directly yields rate constants for individual steps of the binding process while time-resolved experiments can measure rate constants for only the *overall* binding process. In particular, the simulation revealed a two-step binding process in which the formation of a metastable encounter complex intermediate is rate-limiting followed by rearrangement of this nonnative encounter complex to the native complex. Furthermore, sufficient sampling was achieved to calculate the percentage of productive collisions with only 11 ± 5% of all diffusional collisions being productive, *i.e.* eventually resulting in the bound state.

Third, our simulation identified the most kinetically important interactions for the binding process. These interactions, which involve barnase residues, R59bn and S38bn, fasten opposite ends of the binding interface prior to rearrangement of the encounter complex to the native complex.

Finally, short-timescale solvent dynamics during the binding process were resolved in our simulation. In particular, desolvation of the protein binding interfaces was found to occur during the late stage in the binding process just prior to rearrangement of the encounter complex to the native complex. Furthermore, once the bound state is reached, all but one of the nine positions of interfacial crystallographic water molecules that bridge hydrogen bonds between barnase and barstar were occupied by water molecules that originated from the bulk solvent.^26^

Taken together, our WE simulation provide direct views of pathways at an unprecedented level of detail as well as the necessary extensive sampling to validate current simulation models, especially in the accurate calculation of kinetics observables. Given that the simulation would now be completed within 10 days on a GPU using 16 NVIDIA Tesla P100 GPUs at a time, WE-enabled atomistic simulations of multi-μs protein binding processes are now practical on typical resources.

## Acknowledgements

This work was supported by NSF CAREER Award MCB-0845216 and NIH 1R01GM115805-01 to LT Chong and a DAAD graduate research grant to AS Saglam. Computational resources were provided by NSF XSEDE allocation TG-MCB100109 to LT Chong, NSF CNS-1229064, and the University of Pittsburgh’s Center for Research Computing. We thank Alex DeGrave and Alex Dickson for providing plotting tools; and Daniel Zuckerman (OHSU), Terrance Oas (Duke University), AJ Pratt, and Anthony Bogetti for helpful discussions.

